# Real-time Continuous Hand Motion Myoelectric Decoding by Automated Data Labeling

**DOI:** 10.1101/801985

**Authors:** Xuhui Hu, Hong Zeng, Dapeng Chen, Jiahang Zhu, Aiguo Song

## Abstract

In this paper an automated data labeling (ADL) neural network was proposed to streamline dataset collecting for real-time predicting the continuous motion of hand and wrist, these gestures are only decoded from a surface electromyography (sEMG) array of eight channels. Unlike collecting both the bio-signals and hand motion signals as samples and labels in supervised learning, this algorithm only collects the unlabeled sEMG into an unsupervised neural network, in which the hand motion labels are auto-generated. The coefficient of determination (r2) for three DOFs, i.e. wrist flex/extension, wrist pro/supination, hand open/close, was 0.86, and 0.87 respectively. The comparison between real motion labels and auto-generated labels shows that the latter has earlier response than former. The results of Fitts’ law test indicate that ADL has capability of controlling multi-DOFs simultaneously even though the training set only contains sEMG data from single DOF gesture. Moreover, no more hand motion measurement needed which greatly helps upper-limb amputee imagine the gesture of residual limb to control a dexterous prosthesis.

## I. INTRODUCTION

Gesture recognition is one of the most significant research contents in Human-Machine Interaction(HMI). In order to obtain the hand motion information, state of the art technology utilizes a direct measurement method typified by data glove and exoskeleton device^[1]^, which can obtain accurate motion information. However, long-wearing is not comfortable enough, as well as poor adaptability to different hand shapes. In addition, an indirect measurement method typified by computer vision has excellent recognition performance with the help of deep learning^[2]^, but the recognition accuracy is affected by intensity of surrounding light, image background, and obstacle occlusion. Recently, bio-signal based hand recognition has become a hotspot^[3-5]^, which has the advantages of comfortable wearing, unrestricted hand shape and outdoor usage. Since the motion of the hand and wrist is mainly innervated by the forearm muscles, one of the mainstream research directions is to extract potential motion intentions from the forearm muscles signals. Lauren develops a real-time simultaneous hand gesture recognition using intramuscular EMG^[6,7]^. Anvaripour investigates the forearm muscles movement data processing sensed by an array of Force Sensor Resistor^[8]^. Zhang utilizes electrical impedance tomography to image the muscles that govern the gestures and uses image processing algorithms to identify multiple gestures^[9]^. Additional, many researchers are making efforts on analyzing sEMG signals^[10-13]^. To summarize, most of the above studies are based on pattern recognition (PR). PR can recognize stable and discrete gestures during the movement of the hand, but cannot predict the transition state between gestures, which missing the sense of proprioception^[14]^. Currently, machine learning algorithms have been used to carry out the research of continuous motion intention recognition. Jiang designs a proportional and simultaneous controlled prosthetic hand by a dedicated multi-layer perceptron networks^[15]^. Yang proposes a convolutional neural network structure to decode simultaneous movements with three DOFs^[16]^. However, most of the research focuses on supervised learning, that is, while collecting myoelectric signals, it has to collect hand motion signals by other sensors such as camera or data glove at the same time, which increases the difficulty of collecting training samples.

In order to extract potential motion intention from sEMG signal, a large number of studies have proved the theory of muscle synergy^[17]^, that the low-dimensional motion instruction is coordinated by the high-dimensional muscle groups. In addition, since the sEMG signal is the superposition of the motion unit action potential sequence (MUAPs), a large number of MUAPs decoding algorithms use the idea of blind source separation^[18]^, that the low-density motion unit is decoded by the high-density EMG signals. It’s obviously shown that both EMG signal processing tasks use the same dimensionality reduction strategy to extract low-dimensional potential motion intention features from redundant EMG signals. Inspired by the dimension reduction method based on the biomechanical theory, researchers have used various algorithms such as principal component analysis (PCA)^[19]^, non-negative matrix factorization (NMF)^[20,21]^, autoencoders (AEN)^[22]^ to study the continuous motion intention. However, above methods rely entirely on the self-learning of machine, it is difficult to learn a regression model that exactly meets the designer’s expectations when without any expert experience.

In this paper, a myoelectric regression model based on hybrid deep neural network was designed to decode low-dimensional hand motion from high-dimensional sEMG signals. Compared with the traditional supervised learning model, the proposed HDL only rely on the unlabeled sEMG data to learn the gesture regression model. HDL network comprises three layers, the first layer is spatial-temporal feature (STF) layer, which extracts temporally and spatially related high-dimensional principal component features from redundant sEMG signals. The second layer is muscle synergy feature(MSF) layer that uses AEN to learn low-dimensional muscle synergy features. The third layer is the mapping layer of the muscle synergy feature to the actual motion intent. This layer is actually a regression neural network layer, the motion label used by this layer is extracted from MSF layer.

## II. METHOD

### A. Hybrid Deep *Learning Network Framework*

The advantage of deep learning is the capability of automatically fitting an accurate mathematical model by gradient descent algorithm, but this self-learning is random, so it is difficult to assign specific functions to each layer in a deep neural network, which increases the network training duration and the difficulty of algorithm debugging. More seriously, the deeper the network, the harder it is to train and debug. Combining the above advantages and disadvantages, this paper added expert experience to each neural network layer, as shown in Figure 1. The raw sEMG of eight channels were firstly undergoing pre-processing, the means of root mean square (RMS) was adopted to obtain an activation signal. Then these eight activation signals were segmented by a fixed length time window and as input layer of the neural network. As a result, each input sample would contain both spatial and temporal information of array sEMG. The first hidden layer of the network used PCA to reduce the dimension of the input signal. The second hidden layer used AEN to learn the six muscle synergies so as to further reduce the dimensions to neurons. The last hidden layer combined the muscle synergy feature with the auto-generated label from MSF layer into a regression model. Finally, the output layer contained three neurons that present continuous motion data in three DOFs. In summary, the weight matrix of each neural network hidden layer was independently trained and then stacked together, so it can be finely tuned layer by layer during the debugging process.

**Figure 1.**
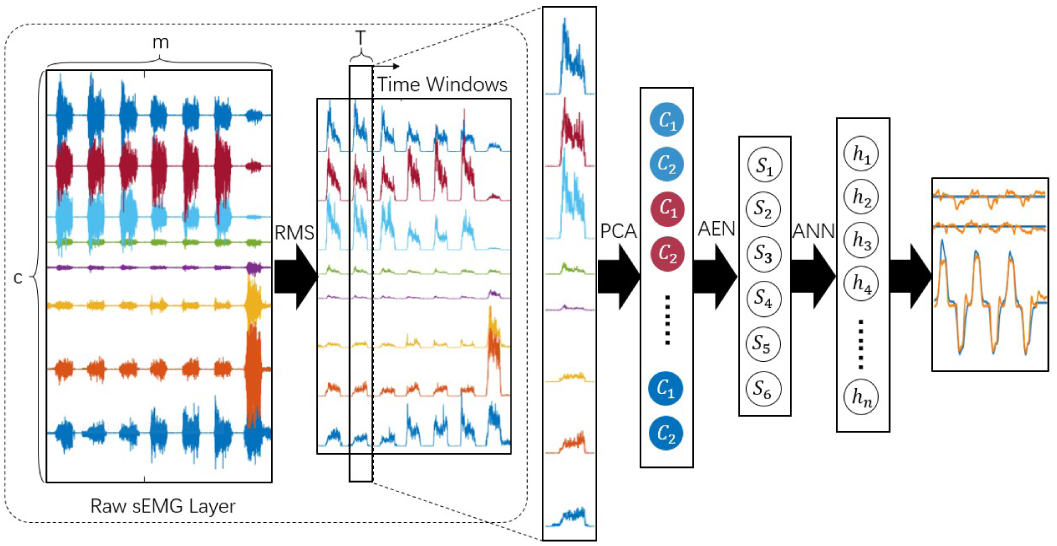
Hybrid Deep Learning Network Framework.

### B. Spatial-Temporal Feature Layer

The sampling rate of sEMG sensor is generally higher than 100Hz, thus the number of input layer neuron will be very large, which is not conducive to network learning. In Figure 1, suppose there are T sample points in each time window and C channels in the sEMG array, so the number of input layer neuron is C×T. In order to obtain representative time and space components among the redundant information, PCA is adopted for temporal analysis on each individual sEMG channel. The PCA input features of a certain sEMG channel are T activation signal sampling points in the time window. The dataset for PCA is generated in the form of a sliding window (suppose the length of sliding step is one, the length of sEMG training dataset on one channel is m, then the number of generated samples is m-T+1). In Figure 1, time series signal of each sEMG channel are compressed into two components.

### C. Muscle Synergy Feature Layer

After processing through the first layer network, the dimension of neurons had reduced from C×T to C×2. Further, inspired by the theory of muscle synergy, the second hidden layer was used to extract six non-negative muscle synergy features (MSF), respectively corresponding to wrist flex/extension, wrist pro/supination and hand open/close. It should be noted that although the six actions described above belong to three DOFs, but in according to the theory of muscle synergy, the two actions of the same DOF still belong to different synergies.

The weight matrix of MSF layer is learned by AEN, as shown in Figure 2. One of its most significant feature is that the input neurons are exactly the same as the output neurons, and the number of hidden layer neurons is smaller than the neurons at both ends of AEN. Therefore, this unique structure can obtain some potential features from the input information. In Figure 2, the network from input layer to hidden layer is called encoding process (circled with a blue box), from the hidden layer to the output layer is called decoding process, we use encoding part as MSF layer. In order to obtain non-negative hidden layer neurons, the Relu function is adopted as the activation function in the encoding process. Since the input layer neurons is calculated by PCA that contains negative components, in order to recover negative features of the output neurons, the Tanh Function is adopted as activation function in the decoding process. A cross entropy function was used as the loss function of AEN; the weight matrix of AEN was initialized using Xavier method; Dropout method was adopted during the iterative training process to prevent over-fitting; the training speed was accelerated by ADAM method and Mini-Batch method. Compared with the existing AEN based myoelectric regression model, we used AEN as the hidden layer of the network, which mainly provided the basis for the extraction of motion intention labels, not for directly learning the final gesture.

**Figure 2.**
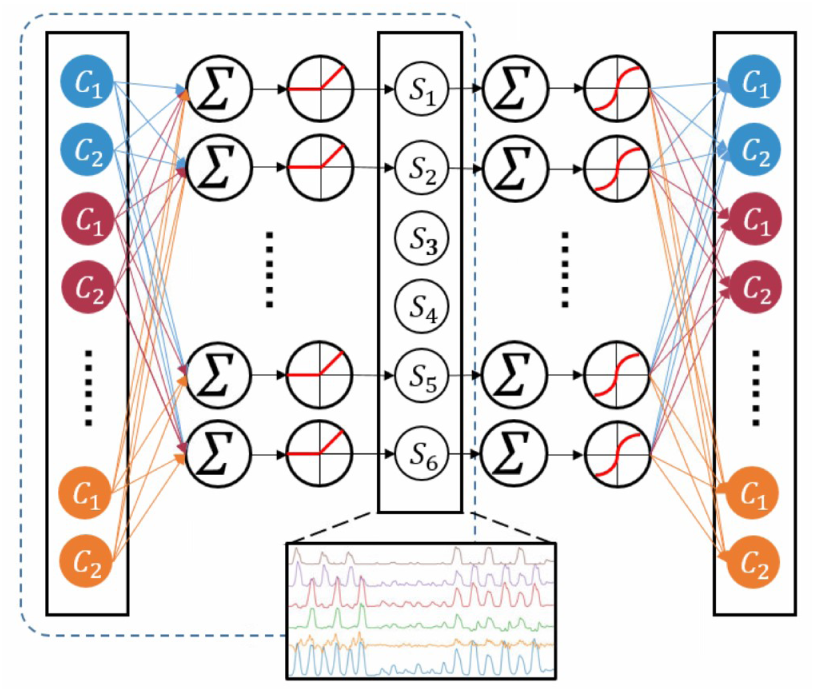
AEN Layer Framework.

### D. Regression Neural Network Layer

Although the muscle synergies learned from AEN are subject to some expert experience, the randomness inherent in the gradient descent will make it difficult to extract the desired motion intention directly. Some research made effort on adding the independence constraints of different neurons, or increasing the training round to screen out the ideal training results, but consequential problems are longer training and selecting time.

Even though the six neurons originally designed in MSF layer may not be able to learn the six motion intentions as expected, our subsequent experiments showed that real intentions are just coupled with each other and form these features. However, when there is no label, the neural network lacks an intuitive learning direction to decouple them.

Here presenting a method for extracting potential motion intentions from MSF. As the six features are subjected to the vector superposition operation, we get a blue oscillating waveform series shown in Figure 3. Each peak indicates the maximum activation of muscle synergy at a certain action; Each valley indicates the resting state of muscle. Therefore, a complete valley-peak-valley segment represents a motion procedure from the beginning of the hand gesture to the strongest muscle activation and restore to rest again. In this paper, by searching for peaks and valleys, the algorithm can automatically reconstruct a motion intention label with three DOFs for the hand and wrist.

**Figure 3.**
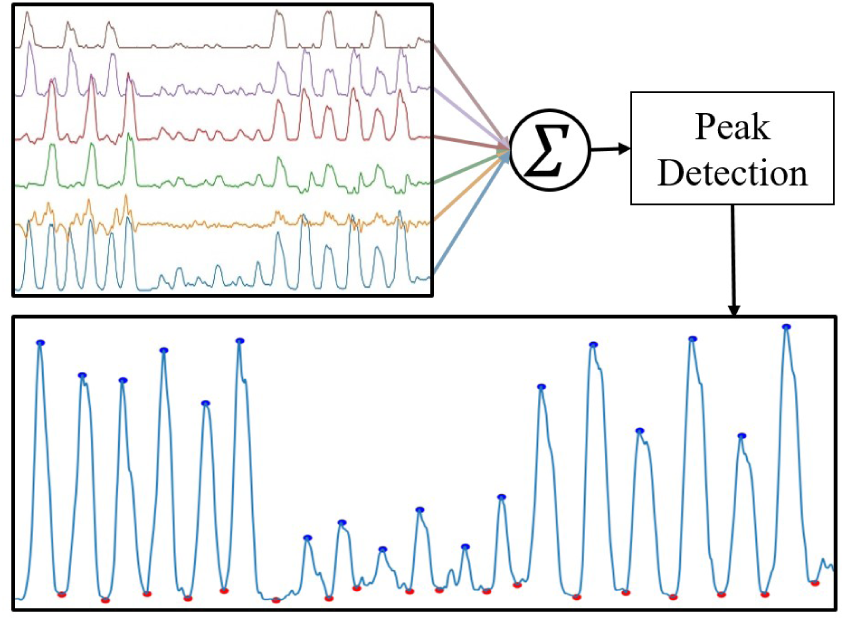
Auto-Generated Motion Label Flow Chart.

In Figure3, the blue waveform depicts a round of data collection procedure, the gesture training sequence is wrist flex/extension three times; wrist pro/supination three times; hand open/close three times. It’s obvious that the muscle activation level of each gesture is different, the reason is that the muscle innervated for wrist rotation is located inside the deeper forearm muscle group, and these muscles are smaller, as a result, the acquired muscle activation is lower than other gestures. Therefore, it is necessary to perform uniform normalization processing on different gestures. After normalization, the maximum activation of each gesture is set to one, and the rest status is zero. After obtaining normalized the hand gesture label, a regression neural network (RegNN) is built for mapping the MSF layer to the label, then the trained regression network layer is stacked with the STF layer and MSF layer to form the final DL model.

Comparing with the method of data glove to acquire hand gesture, this paper utilizes sEMG signal to achieve continuous motion intention. Although latter has lower accuracy than previous, but has quicker response because nerve electrical signal conductive velocity is faster than actual muscle movement velocity.

## III. EXPERIMENTAL PARADIGM

### A. Subjects and Experimental Platform

Six able-bodied subjects (24±2 years old) were enrolled in real-time recognition experiment. The experiment complied with the Helsinki Declaration. All subjects signed written informed consent, and the Ethics Committee of Southeast University approved the protocol.

In the experiment, all subjects wore a Thalmic Lab’s MYO armband to collect sEMG signals. The sensor consists of eight equidistant, independent channels of bipolar electrodes with reference electrodes at the middle of each group of electrodes, eight reference electrodes are conductive to each other. The advantage of MYO is easy to wear and adaptable to users’ arm circumferences. The sampling frequency of the sensor is 200Hz, the AD precision is 8 bits.

The collected sEMG signals were sent to a laptop via Bluetooth communication, using Python 3.6 as programming language. STF layer was implemented by scikit’s PCA library, and MSF layer was implemented by tensorflow framework. The regression layer was implemented by scikit’s regression neural network library. The hardware for training DL network was intel i5 7300HQ at 2.5GHz.

### B. Dataset Collection Procedure

Before starting the training, the experimental subjects were firstly informed about the relevant experimental content to get familiar with the experimental movements. The subjects were required to make six gestures (wrist flex/extension, wrist pro/supination and hand open/close) followed by a virtual hand as shown in Figure 4, the same gesture was made three times in a row, then the next gesture. Each movement lasted for 3 seconds from the beginning to the completion, and then the hand was in a relaxed state. When all the gestures were done, it was regarded as the end of one round of collection.

**Figure 4.**
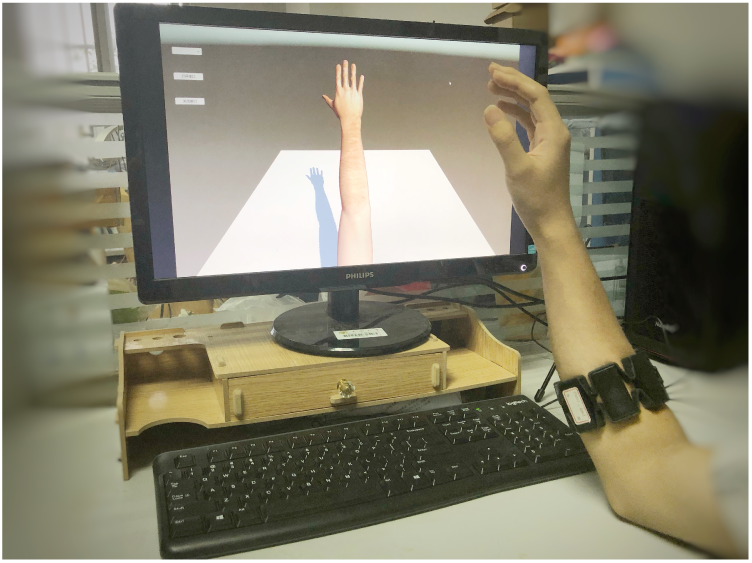
Experimental Platform of Data Collection and Test.

**Figure 5.**
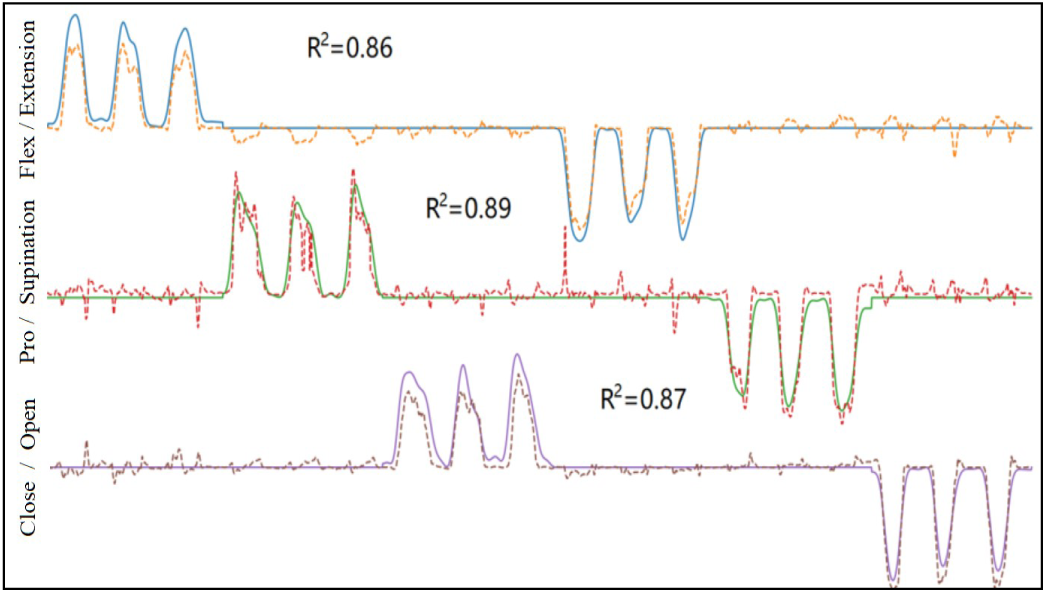
Fitting curve on test set.

Each subject conducted three rounds of data collection procedure, the first two rounds were utilized to train the deep neural network, the last round was seen as test set to validate the offline performance of algorithm. When the network is trained completely, subject can control the virtual hand in real time to show the performance of algorithm

## IV. RESULTS

### A. Offline Analysis

In this section, an offline analysis of sEMG test set was performed to illustrate the decoding performance of the DL algorithm. Since the core of the proposed method was data compression, Figure 6 shows the results before and after compression in each neural layer of DL network, as well as the auto-generated labels (AGL) extracted in MSF layer.

**Figure 6.**
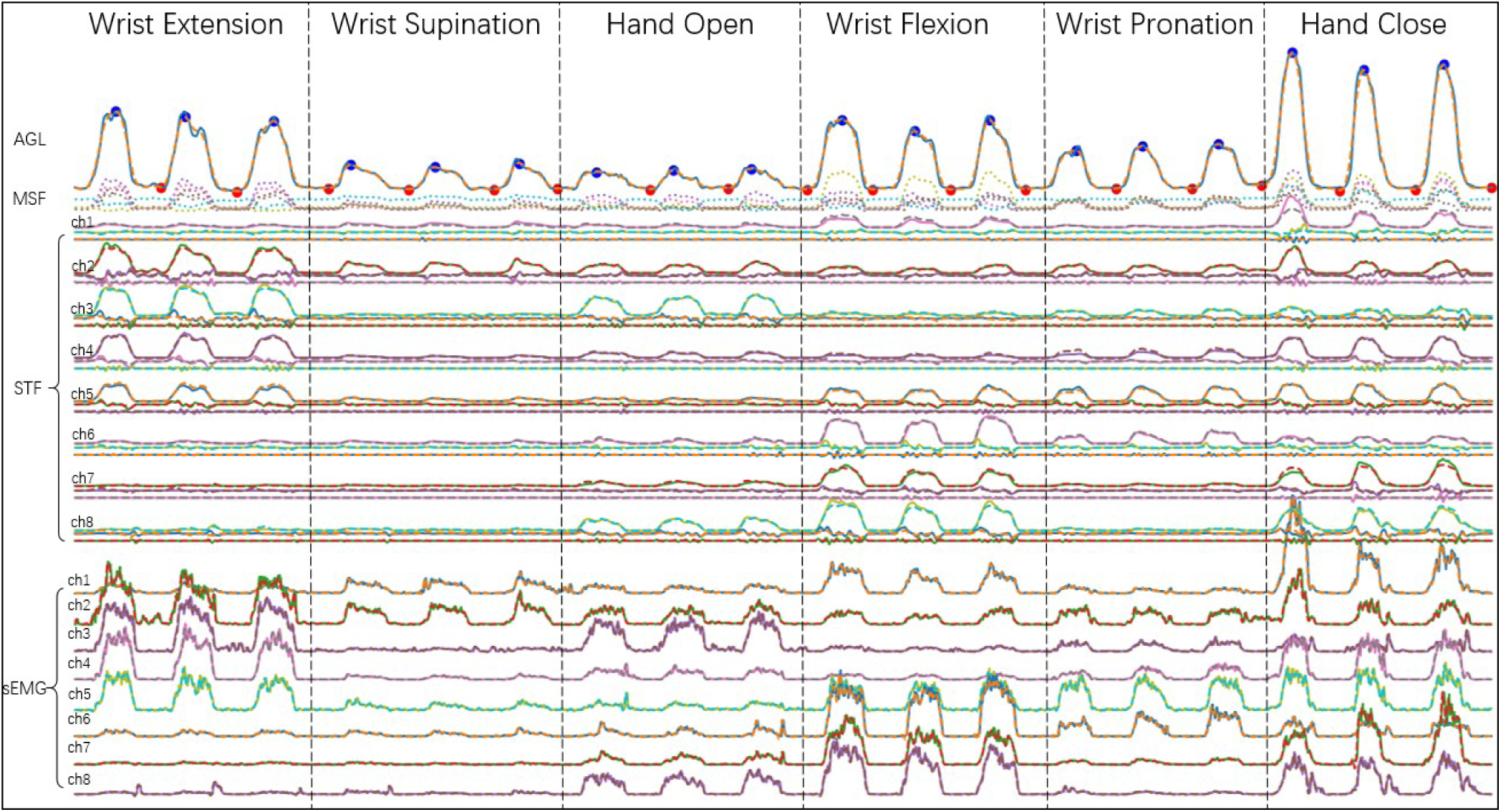
Compression, Reconstruction and Extraction in the test set at all layers of DL algorithm.

From bottom to up of Figure 6, 8 channels of RMS sEMG are firstly compressed by the STF layer, where the dotted line is reconstruction data, the average r2 of eigh channels is 0.979. Secondly, the first three principal components of each sEMG channel were extracted from the preliminary experiment. 24 dimensions of STF are compressed by the MSF layer, the average r2 of the first principal component among all channels is 0.962, the second is 0.530, and the third is -0.007. It can be seen that MSF layer is basically unable to reconstruct the third component, so the neurons of the actual STF layer can be compressed to 16.

After two-stage compression, 160 features of temporal and spatial related sEMG input are finally compressed to 6-dimensional muscle synergy feature, which are drawn above STF by dotted lines.

In order to extract AGL, the vector superposition of six MSFs was calculated and showed on the top of Figure 6, the peak and valley of AGL was found using extremum searching algorithm. Finally, three dimensional label can be identified according to the sequence order of gestures in the data collection phase, as shown by solid line in Figure 5. Since the muscle activation of different gestures varies, it is necessary to perform a standard normalization process on the intervals of different gestures. In addition, in order to improve the fitting effect, a bidirectional Butterworth digital filter was used to obtain a smooth gesture label. In this paper, low-pass filtering with a cutoff frequency of 1.65 Hz can obtain label data with better response and smoothness. Figure 5 shows the fitting curve of test set in the RegNN layer, the solid line is gesture label, and the dotted line is the actual predicted gesture. The r2 of three gestures are 0.86, 0.89, and 0.87.

There is another method of obtaining AGL, that is directly superimposes RMS sEMG of eight channels. We compared this method (represented as RMS) with the proposed MSF method, and referred to the actual motion parameters collected by Leap Motion (represented as LEAP). In Figure 7, curves in the upper figure is actual label representing one subject repeated three times of hand close, the labels of three different methods were independently standard normalized. Curves in the bottom figure are the one-order derivative of above curves, it shows that the response of MSF has 113±26 ms faster than that of RMS, but 233±34ms slower than that of LEAP during the beginning of gesture, while MSF has fastest response during the end of gesture,that is 106±47 ms faster than RMS and 533±38 ms faster than LEAP. As a result, MSF has better response than RMS, even better than actual motion label during the end of the gesture.

**Figure 7.**
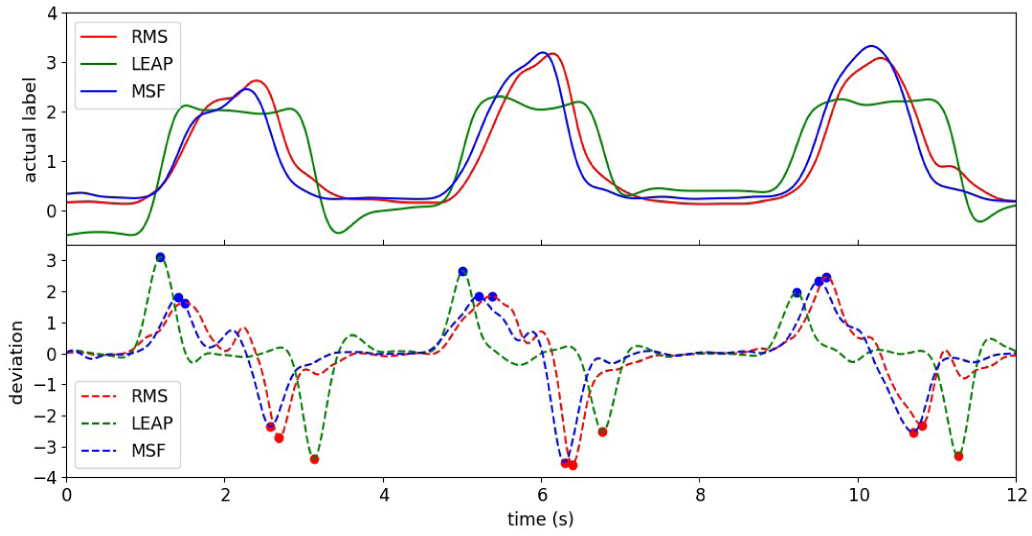
Response Test between RMS,MSF and LEAP.

**Figure 8.**
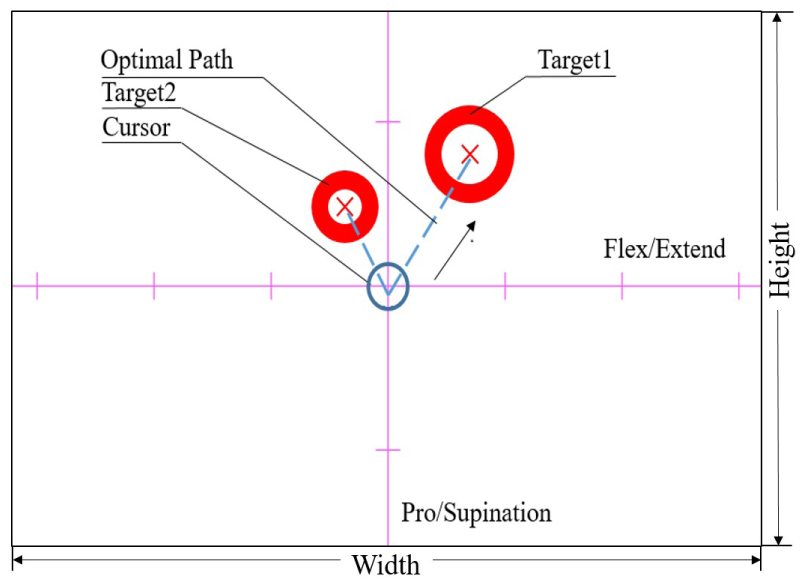
Virtual Test Scenario Based on Fitts’ Law.

### B. Online Test

Since Fitts’ Law testing has been extensively used in the validation of HMI^[23]^, an 2D evaluation scenario based on Fitts’ law test, as shown in Figure 8, was established to quantify the online simultaneous performance of DL algorithm.

In this experiment, subjects were required to move the cursor (blue circle) to the target (red circle). Two alternative control strategies exist in different researches^[24,25]^ were compared. In the first control strategy, the wrist flex/extension controlled the horizontal movement of the cursor, the wrist rotation (pro/supination) controlled the vertical movement of the cursor (represented as ROT); Another strategy used the same horizontal control strategy, but utilized radial/ulnar deviation to control the vertical movement (represented as DEV).

The resolution of the virtual test scenario was 800×600 pixels(Width×Height). Each subject needed to complete 20 times of hitting target tasks, the size and location of the target point were randomly generated. There was only one target point in the virtual scene at a time. When the cursor successfully hitting the target, the original target point disappeared and next target point appeared.

The speed of the cursor was controlled by the output of the DL network, and the actual speed mapped to the virtual scene was determined empirically. The movement speed finally up to 450 pixels per second by increasing the speed to maximize the control effect. Each task had a 15s countdown. If timed up, the task would fail. When the distance between cursor and target was smaller than the radius of target, the cursor needed dwelling time of 1s. If the cursor passed through the target before the dwelling time was reached, this task would be marked as overshoot.

In Fitts’ Law Testing, random targets in a virtual scene are labeled with index of difficulty (ID) based on the distance (D) between target and cursor origin and the radius (R) of target, as shown in the Formula 1. In this experiment, R was set as 50 to prevent the ID of targets in the scenario from exceeding 4.

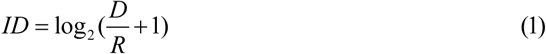

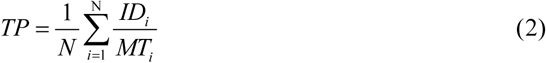

Five performance metrics were selected in this experiment, respectively completion rate (CR), movement time (MT), overshoot (OV), throughput (TP) and path efficiency (PE). Table II demonstrates the meaning of specific metric.

**TABLE I.** COEFFICIENT OF DETERMINATION ALONE NEURAL LAYERS

|  | sEMG | 1<sup>st</sup> component | 2<sup>nd</sup> component | 3<sup>th</sup> component |
| --- | --- | --- | --- | --- |
| 1 | 0.980 | 0.838 | 0.531 | -0.004 |
| 2 | 0.969 | 0.990 | 0.642 | -0.016 |
| 3 | 0.982 | 0.990 | 0.383 | -0.007 |
| 4 | 0.981 | 0.977 | 0.600 | -0.008 |
| 5 | 0.973 | 0.975 | 0.719 | -0.006 |
| 6 | 0.984 | 0.988 | 0.391 | -0.012 |
| 7 | 0.982 | 0.958 | 0.481 | -0.004 |
| 8 | 0.983 | 0.980 | 0.495 | -0.001 |

**TABLE II.** METRICS IN FITTS’ LAW EXPERIMENT

| Metric | Description |
| --- | --- |
| CR(%) | The ratio of completed tasks to all tasks. |
| MT(s) | The time it took the subject to complete the task of corresponding ID. |
| OV(%) | The ratio of overshoot tasks to all tasks |
| TP(bit/s) | the average ratio of ID of each target to corresponding MT, as shown in Formula 2. |
| PE(%) | the ratio between the length of the optimal path from the initial point to the target and the actual trajectory realized |

**TABLE III.** RESULTS OF THE TWO DIMENSIONAL FITTS’ LAW TEST

| Metrics | ROT | DEV |
| --- | --- | --- |
| CR(%) | 100.00 | 100.00 |
| MT(s) | $4.99 \pm 2.33$ | $3.01 \pm 0.97$ |
| OV(%) | $21 \pm 6$ | 0.00 |
| TP(bit/s) | $0.55 \pm 0.19$ | $0.87 \pm 0.18$ |
| PE(%) | $56 \pm 17$ | $78 \pm 11$ |

In Figure 9, a stronger linear relation is found between CT and ID in DEV (r2=0.66) than in ROT (r2=0.39), it indicates that DEV are much suitable of applying the Fitts’ Law test.

**Figure 9.**
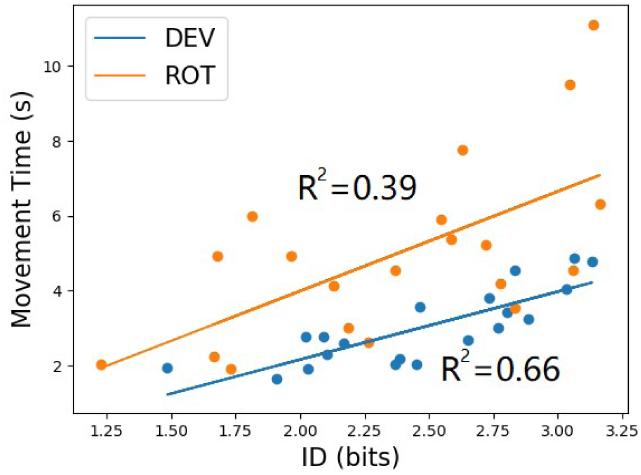
linear relation between CT and ID.

Two control strategies were compared on actual tracking paths. In Figure 10, it shows that more sharp turns are on the path of ROT, indicating poorer performance on simultaneous control. By contrast, DEV can better track targets located in non-axis directions. There are two major reasons for these difference results. On the one hand, rotating and flex/ extending the wrist simultaneously has weak proprioception on the subject’s sense of direction control. On the other hand, because of the lower activation of the wrist muscle, subjects has to rotate their wrist to larger angles to control cursor movement. Meanwhile, wrist flex/extension is more sensitive than rotation, making it difficult to synchronize two gestures. The DEV is more in line with the intuitive control intention of the directions, and the signals of radial/ulnar deviation are more synchronized, which makes it easier to predict two gestures at the same time. In this paper, the same algorithm is used to identify the gesture of the DEV strategy. The r2 of three gestures are 0.90 (wrist flex/extension), 0.89(wrist deviation), 0.86 (hand open/close).

**Figure 10.**
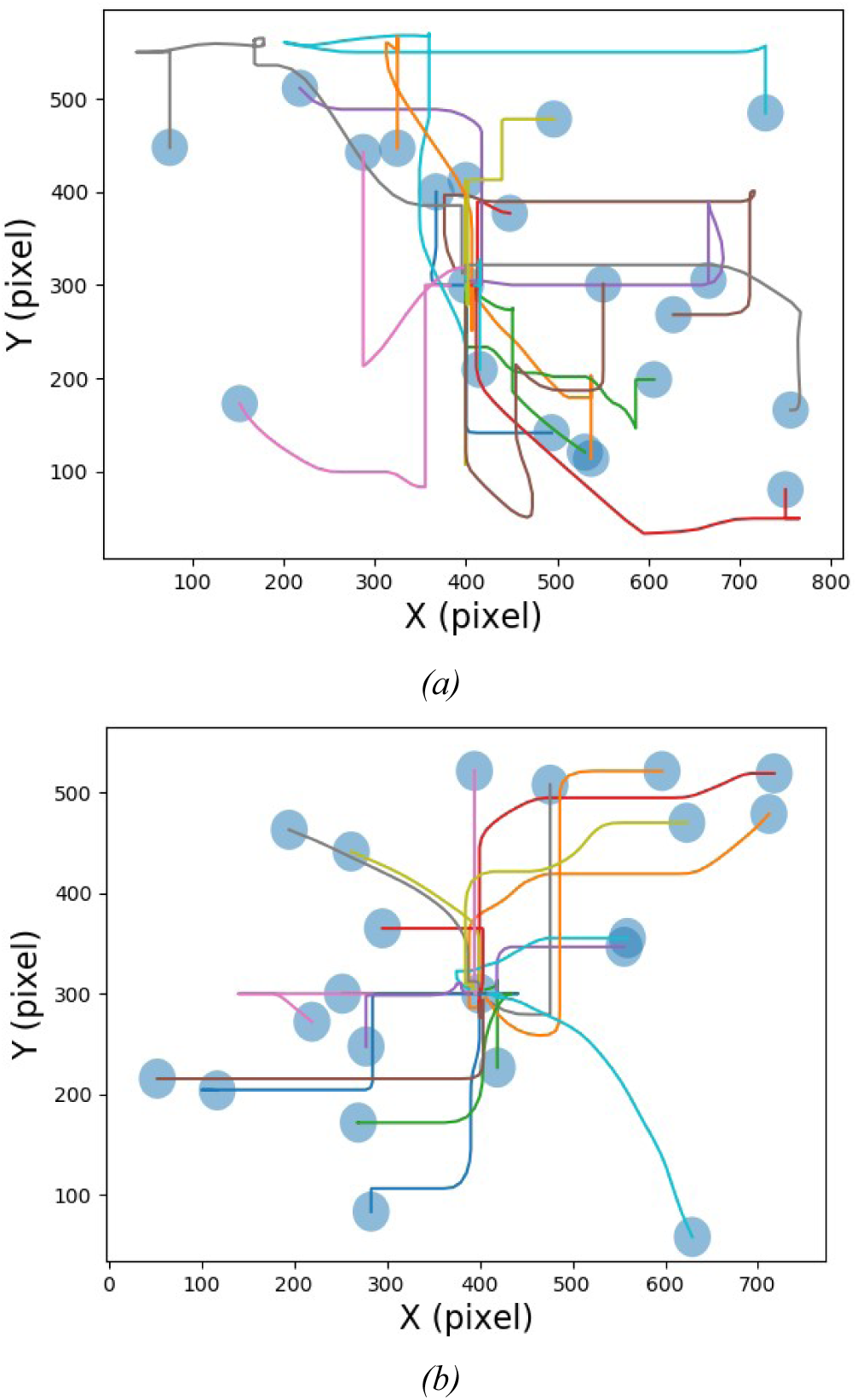
Hitting Target Path of ROT (a) and DEV (b)

Table III compares five metrics of DEV and ROT. It shows that both strategies can complete the task within the specified time, but DEV is better than ROT on the completion effect of tasks.

More importantly, both strategies proves that Even if the training dataset does not include the simultaneous gestures (a gesture that combines from two different DOFs), the algorithm can still achieve correct predictions when actual simultaneous gestures occurred, which also approves that muscle synergy feature from single DOF gesture can represent multi-DOFs gesture.

## V. CONCLUSIONS AND FUTURE WORK

This paper proposes a HDL algorithm for real-time predicting the simultaneous hand gestures of three DOFs, which are only decoded from a sEMG array of eight channels. HDL collects the unlabeled sEMG to auto-generate the potential motion label. The comparison between real motion label and auto-generated label shows that the latter has earlier response than former. The results of Fitts’ law test indicate that HDL has capability of controlling multi-DOFs simultaneously even though the training set of HDL only contains sEMG data from single DOF gesture. Moreover, No more hand motion measurement needed which greatly helps upper limb amputee learn the gesture of residual limb to control a dexterous prosthesis.

There are several additional avenues of future work. On the one hand of HMI, HDL shortens the training time with the help of layer-by-layer training and expert experience, but further research on the generalization performance is necessary to make the algorithm adapt to different individuals quickly^[26,27]^. On the other hand of sEMG controlled prosthesis, we conducted the preliminary test on upper limb amputee and found that the recognition sharply deteriorated before and after wearing the prosthetic hand, which is a difficult problem in the field of prosthetic research^[28,29]^. Therefore, we will further carry on how to improve the robustness of prosthetic control.

## REFERENCES

[1] Neto P, Pereira D, Pires J N, et al. Real-time and continuous hand gesture spotting: An approach based on artificial neural networks[C]. 2013 IEEE International Conference on Robotics and Automation, 2013: 178–183.

[2] Ren Z, Yuan J, Zhang Z. Robust hand gesture recognition based on finger-earth mover’s distance with a commodity depth camera[C]. Proceedings of the 19th ACM international conference on Multimedia, 2011: 1093–1096.

[3] Agashe H A, Contreras-Vidal J L. Decoding the evolving grasping gesture from electroencephalographic (EEG) activity[C]. 2013 35th Annual International Conference of the IEEE Engineering in Medicine and Biology Society (EMBC), 2013: 5590–5593.

[4] Zeng H, Wang Y, Wu C, et al. Closed-Loop Hybrid Gaze Brain-Machine Interface Based Robotic Arm Control with Augmented Reality Feedback[J]. Frontiers in Neurorobotics, 2017, 11(60).

[5] Wu C, Zeng H, Song A, et al. Grip Force and 3D Push-Pull Force Estimation Based on sEMG and GRNN[J]. Frontiers in Neuroscience, 2017, 11(343).

[6] Smith L H, Kuiken T A, Hargrove L J. Real-time simultaneous and proportional myoelectric control using intramuscular EMG[J]. Journal of Neural Engineering, 2014, 11(6): 066013.

[7] Smith L H, Kuiken T A, Hargrove L J. Evaluation of Linear Regression Simultaneous Myoelectric Control Using Intramuscular EMG[J]. IEEE Trans Biomed Eng, 2016, 63(4): 737–46.

[8] Anvaripour M, Saif M. Hand gesture recognition using force myography of the forearm activities and optimized features[C]. 2018 IEEE International Conference on Industrial Technology (ICIT), 2018: 187–192.

[9] Zhang Y X R, Harrison C. Advancing hand gesture recognition with high resolution electrical impedance tomography[C]. Proceedings of the 29th Annual Symposium on User Interface Software and Technology - UIST ‘16, 2016: 843–850.

[10] Geng W, Du Y, Jin W, et al. Gesture recognition by instantaneous surface EMG images[J]. Sci Rep, 2016, 6: 36571.

[11] Amma C, Krings T, Böer J, et al. Advancing Muscle-Computer Interfaces with High-Density Electromyography[C]. Proceedings of the 33rd Annual ACM Conference on Human Factors in Computing Systems - CHI ‘15, 2015: 929–938.

[12] Daohui Zhang X Z, Jianda Han, and Yiwen Zhao. A Comparative Study on PCA and LDA Based EMG Pattern Recognition for Anthropomorphic Robotic Hand*[C]. ICRA, 2014.

[13] Yang D, Yang W, Huang Q, et al. Classification of Multiple Finger Motions During Dynamic Upper Limb Movements[J]. IEEE Journal of Biomedical and Health Informatics, 2017, 21: 134–141.

[14] Farina D, Jiang N, Rehbaum H, et al. The extraction of neural information from the surface EMG for the control of upper-limb prostheses: emerging avenues and challenges[J]. IEEE Trans Neural Syst Rehabil Eng, 2014, 22(4): 797–809.

[15] Jiang N, Vest-Nielsen J L G, Muceli S, et al. EMG-based simultaneous and proportional estimation of wrist/hand kinematics in uni-lateral trans-radial amputees[J]. Journal of Neuroengineering and Rehabilitation, 2012, 9.

[16] Yang W, Yang D, Liu Y, et al. Decoding Simultaneous Multi-DOF Wrist Movements From Raw EMG Signals Using a Convolutional Neural Network[J]. IEEE Transactions on Human-Machine Systems, 2019: 1–10.

[17] D’avella A, Saltiel P, Bizzi E. Combinations of muscle synergies in the construction of a natural motor behavior[J]. Nature Neuroscience, 2003, 6(3): 300–308.

[18] Farina D, Holobar A. Human? Machine Interfacing by Decoding the Surface Electromyogram [Life Sciences][J]. IEEE Signal Processing Magazine, 2015, 32(1): 115–120.

[19] Hargrove L J, Li G, Englehart K B, et al. Principal components analysis preprocessing for improved classification accuracies in pattern-recognition-based myoelectric control[J]. IEEE Trans Biomed Eng, 2009, 56(5): 1407–14.

[20] Jiang N, *, Englehart K B, et al. Extracting Simultaneous and Proportional Neural Control Information for Multiple-DOF Prostheses From the Surface Electromyographic Signal[J]. IEEE Transactions on Biomedical Engineering, 2009, 56(4): 1070–1080.

[21] Lin C, Wang B, Jiang N, et al. Robust extraction of basis functions for simultaneous and proportional myoelectric control via sparse non-negative matrix factorization[J]. J Neural Eng, 2018, 15(2): 026017.

[22] Vujaklija I, Shalchyan V, Kamavuako E N, et al. Online mapping of EMG signals into kinematics by autoencoding[J]. J Neuroeng Rehabil, 2018, 15(1): 21.

[23] Scheme E J, Englehart K B. Validation of a selective ensemble-based classification scheme for myoelectric control using a three-dimensional Fitts’ Law test[J]. IEEE Trans Neural Syst Rehabil Eng, 2013, 21(4): 616–23.

[24] Gusman J, Mastinu E, Ortiz-Catalan M. Evaluation of Computer-Based Target Achievement Tests for Myoelectric Control[J]. IEEE J Transl Eng Health Med, 2017, 5: 2100310.

[25] Carles Igual J I, Janne M. Hahne, and Lucas C. Parra. Adaptive Auto-Regressive Proportional Myoelectric Control[J], 2019.

[26] Castellini C, Bongers R M, Nowak M, et al. Upper-Limb Prosthetic Myocontrol: Two Recommendations[J]. Front Neurosci, 2015, 9: 496.

[27] Cote-Allard U, Fall C L, Drouin A, et al. Deep Learning for Electromyographic Hand Gesture Signal Classification Using Transfer Learning[J]. IEEE Trans Neural Syst Rehabil Eng, 2019.

[28] Hahne J M, Schweisfurth M A, Koppe M, et al. Simultaneous control of multiple functions of bionic hand prostheses: Performance and robustness in end users[J]. Science Robotics, 2018, 3(19).

[29] Yang D, Gu Y, Thakor N, et al. Improving the functionality, robustness, and adaptability of myoelectric control for dexterous motion restoration[J]. Experimental Brain Research, 2019, 237.

